# FLECS Technology for High-Throughput Screening of Hypercontractile Cellular Phenotypes in Fibrosis: A Function-First Approach to Anti-Fibrotic Drug Discovery

**DOI:** 10.1101/2023.07.21.549589

**Authors:** Yao Wang, Enrico Cortes, Ricky Huang, Jeremy Wan, Boris Hinz, Robert Damoiseaux, Ivan Pushkarsky

**Affiliations:** Forcyte Biotechnologies, Inc, Los Angeles, CA, 90095; Laboratory of Tissue Repair and Regeneration, Keenan Research Centre for Biomedical Science of the St. Michael’s Hospital, 209 Victoria Street, Toronto, ON M5B 1T8, Canada; Faculty of Dentistry, University of Toronto, Toronto, M5S 3E2 Ontario, Canada; University of California, Los Angeles, Los Angeles, CA, 90095; California NanoSystems Institute at UCLA, Los Angeles, Los Angeles, CA, 90095

## Abstract

The pivotal role of myofibroblast contractility in the pathophysiology of fibrosis is widely recognized, yet HTS approaches are not available to quantify this critically important function in drug discovery. We develop, validate, and scale-up a HTS platform that quantifies contractile function of primary human lung myofibroblasts upon treatment with pro-fibrotic TGF-β1. With the fully automated assay we screened a library of 40,000 novel small molecules in under 80 h of total assay run-time. We identified 42 hit compounds that inhibited the TGF-β1-induced contractile phenotype of myofibroblasts, and enriched for 19 that specifically target myofibroblasts but not phenotypically related smooth muscle cells. Selected hits were validated in an *ex vivo* lung tissue models for their inhibitory effects on fibrotic gene upregulation by TGF-β1. Our results demonstrate that integrating a functional contraction test into the drug screening process is key to identify compounds with targeted and diverse activity as potential anti-fibrotic agents.

## Introduction

Fibrosis is a chronic pathological wound repair process characterized by progressive tissue scarring, resulting in the replacement of normal tissue with thickened and stiffened non-functional scar extracellular matrix (ECM). Central to the disease process that can affect all organs is the activation and maintenance of fibroblasts and other progenitors into myofibroblasts. Myofibroblasts in fibrotic organs exist in a spectrum of activation states that start with the excessive deposition and remodeling of collagen-rich ECM. In the process, myofibroblast develop contractile actin-myosin bundles (stress fibers) that incorporate alpha smooth muscle actin (α-SMA), resulting in a ‘super-contractile’ phenotype. It is the transmission of high contractile force that leads to ECM stiffening and formation of a scar that impedes and often obliterates organ function (Fig. 1). In addition to compromising normal organ function, stiffened ECM perpetuates fibrosis by driving further activation of mechano-sensitive and mechano-responsive myofibroblasts. Another part of this fibrotic feed-forward loop is the mechanical activation of TGF-β1 from latent complexes in the ECM^1,2^ by force transmission via the αv integrins of contracting fibroblastic cells^3,4^. TGF-β1 is a master regulator of fibrosis that, among multiple pro-fibrotic actions, promotes further fibroblast-to-myofibroblast conversion via the canonical Smad2/3 pathway and non-canonical signaling pathways^1,5,6^. Finally, myofibroblast contraction-induced strain may protect collagen ECM from enzymatic degradation^7–9^, all resulting in a net accumulation of ECM and overall stiffer fibrotic tissue.

**Fig.1:**
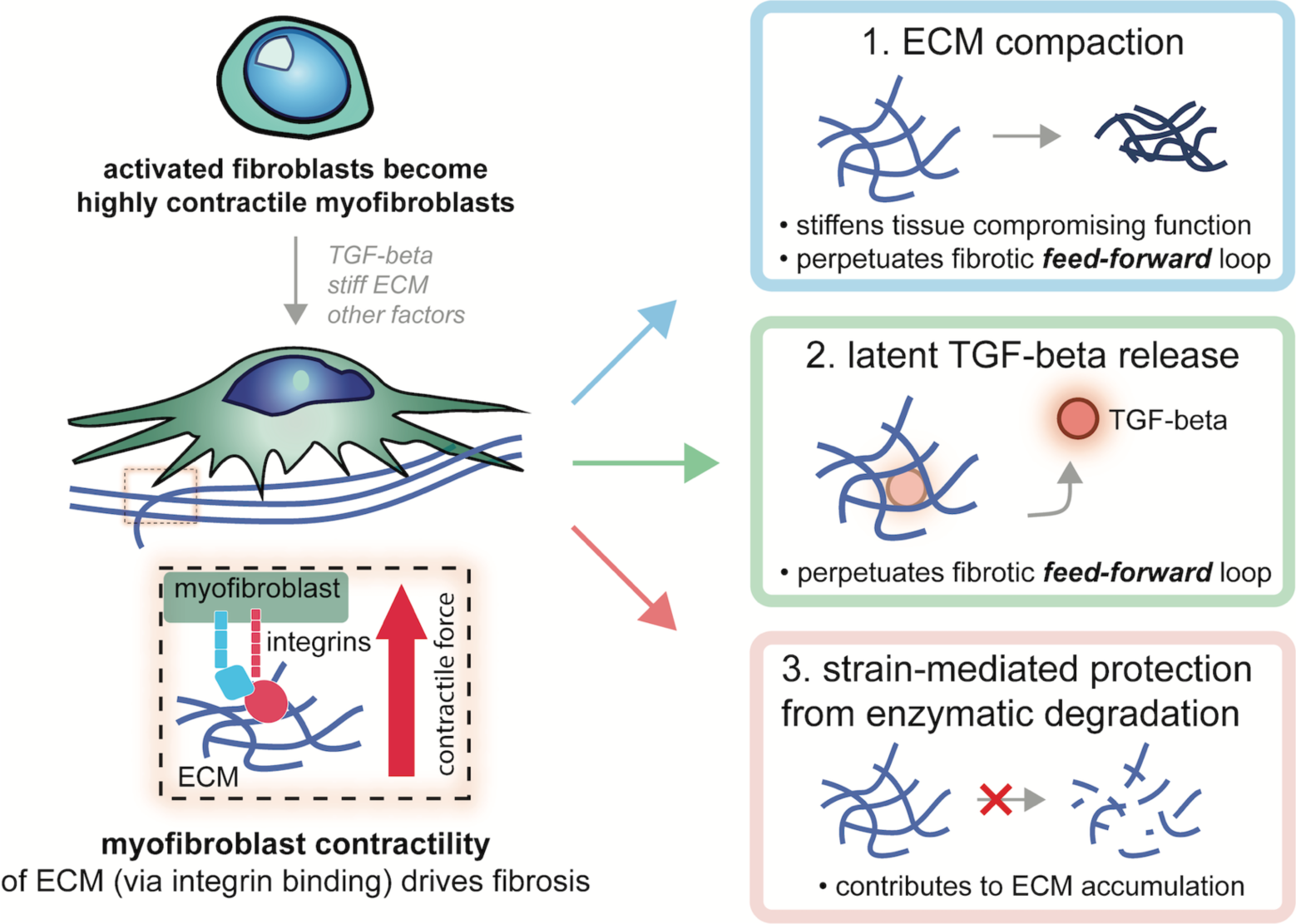
Diagram outlining the three major roles of contractile myofibroblasts in the pathophysiology of fibrosis. (1) The mechanical force exerted on the extracellular matrix (ECM) by myofibroblasts leads to tissue compaction and stiffening, impairing normal function. (2) Myofibroblasts mechanically activate and release Transforming growth factor-beta 1 (TGF-β1) from its latent stores within the ECM, promoting further fibroblast-to-myofibroblast conversion. (3) Concurrently, strain-mediated protection from enzymatic degradation occurs within fibrotic tissue under stress from myofibroblast contraction, resulting in an imbalance between ECM synthesis and degradation, which ultimately leads to net ECM accumulation and tissue thickening.

Despite the widely recognized role of myofibroblast contractility in the pathophysiology of fibrosis, most *in vitro* models of fibrosis, focus primarily on molecular markers of activated myofibroblasts, e.g. expression of the contractile protein alpha-smooth muscle actin (α-SMA) or ECM proteins like pro-collagen, collagen, and fibronectin^10^. Functional assessment of hypercontractile myofibroblasts is often neglected, yet enhanced contraction is a defining feature of myofibroblasts^11,12^. This neglect is partly due to the lack of straight-forward quantitative assays to measure cell contraction. Functional two-dimensional cell growth assays to measure myofibroblast contraction *in vitro* include traction force microscopy (TFM)^10^ and ‘wrinkling’ assays^11^. Both approaches are based on the principle that the contraction of adherent cells results in deformations of planar deformable culture substrates that can be traced through surface marker displacements in TFM or formation of visible large folds (wrinkles). The most widely used three-dimensional myofibroblast contraction assays are based on the diameter reduction of 3D collagen gels by mixed-in fibroblast populations^13,14^, or measurements with directly attached force transducers^12^. Three-dimensional contraction assays cannot deliver information on contraction force of single cells and none of the two-dimensional assays assessing single cell contraction are suitable for higher throughput screening applications. Thus, there is a pressing need for a quantitative and scalable assay that can assess contractility in a fibrosis-relevant context. Such an assay would enable direct screening for novel anti-fibrotic drugs and genes that selectively modulate contractile cell function.

We here develop and validate a functional-phenotypic assay evaluating myofibroblast contractility as a directly observable and quantifiable endpoint, scaled for high-throughput screening (HTS). We built the assay on a previously established high-throughput cell contractility screening platform, that we coined fluorescent elastomeric contractible surfaces (FLECS)^13–19^. In a screening campaign conducted over 15 days, we tested 40,000 small molecules and identified over a dozen inhibitors of myofibroblast contraction with sub-micromolar potency. Our results illustrate the capacity of the My-FLECS assay (FLECS technology applied to screening myofibroblast contractility) to efficiently identify novel starting points for the development of anti-fibrotic drugs within expansive chemical libraries. The inhibitors identified also demonstrated selectivity toward fibroblasts over smooth muscle cells in both the primary assay and in counter-screens, along with the ability to suppress fibrotic gene upregulation in the human precision-cut lung slice model. Collectively, our study underscores the potential of the new screening assay in uncovering novel anti-fibrotic drug activities, which may not be predictable from molecular readouts alone.

## Materials and Methods

### Cell Culture

Cryopreserved human primary lung fibroblasts (HLF) and primary human tracheal smooth muscle (HTSM) cells were purchased from ScienCell. HLF were maintained in Lung Fibroblast Growth Medium (Cell Applications, product 516-500). HTSM cells were maintained in Ham’s F-12 medium supplemented with 10% fetal bovine serum and 1% penicillin/streptomycin. Both cell types were grown in T75 polystyrene flasks at 37°C with 5% C02 content prior to experimentation.

Trypsin-EDTA (0.05%) was used to re-suspend cells at the start of the experiment. For HLF, experiments in the FLECS assay were done in Dulbecco’s Modified Eagle Medium (DMEM) supplemented with 10% FBS and 1% penicillin-streptomycin. For HTSM, experiments were done in the same Ham’s F12 medium as was used for culturing. All experiments were done using cells at passage 3.

### My-FLECS contractility assay general protocol

The My-FLECS contractility assay was performed using an adaptation of the original FLECS protocol described previously^14^ with modifications made specifically to tailor the assay to use with HLF and TGF-β1, for the first time. To initiate the assay, empty 384-well FLECSplates were filled with 25µL of serum-free Dulbecco’s Modified Eagle Medium (DMEM) supplemented with 1% penicillin-streptomycin. HLF cells were detached from their flasks and resuspended at a concentration of 50,000 cells/mL in DMEM/10%FBS medium. TGF-β1 (Peptrotech) was optionally added to this cell suspension at desired concentrations prior to seeding. The cells were then seeded into the pre-filled FLECSplates, with 25µL being added to each well. After seeding, plates were left at RT for 1.5 hrs to allow for uniform settling of cells to the well bottom after which time plates were moved into a 37 C incubator for 24 hrs.

At 24hrs, 10µL of DMEM/10%FBS containing 1µg/mL Hoechst 33342 live nuclear stain was added to each well 30 minutes prior to imaging using a Molecular Devices ImageXpress automated microscope. Image analysis was performed using Forcyte’s proprietary computer vision algorithms as previously described^14^.

It was determined that 70µm sized “X” shaped micropatterns consisting of type IV collagen arrayed over a stiffness of approximately 8 kPa (Forcyte Biotechnologies, product 384-HC4R-QC10) were the optimal assay parameters to support HLF adhesion and contraction both at baseline and following activation with TGF-β1. These parameters were used in all experiments herein.

### Titration of TGF-β1

The optimal concentration of TGF-β1 was determined by testing the contractile response of HLF over a 24hr period against a row-wise 10-step 2-fold dilution series of TGF-β1 starting at 20ng/mL. The experiment was performed as described above with appropriate amounts of TGF-β1 added to each cell suspension prior to seeding into the FLECSplate.

### Pharmacological validation of My-FLECS with ALK-5 inhibitors

ALK-5 inhibitors SD-208, Galunisertib and RepSox (Sigma) were dissolved at 10mM in DMSO and arranged into a 20-step 2-fold serial dilution column-wise in 4 rows each on a 384-well drug source plate. FLECSplates were pre-filled with 25 µL of serum-free DMEM, and then 100 nL of the ALK-5 inhibitor solutions were transferred from their source plate to the corresponding wells on the FLECSplate using a Biomek FXP pin tool. The FLECSplates were shaken at 100 RPM for 15 minutes to ensure mixing. HLF cells were seeded onto the FLECSplates without TGF-β1 and incubated with the ALK-5 inhibitors for 30 minutes. Following incubation, TGF-β1 treatment was performed by adding 10 µL of media containing TGF-β1 to columns 2-23, achieving a final concentration of 2.5 ng/mL, while column 1 received 10 µL of media without TGFβ as the negative control. Plates were imaged after 24hrs as previously described. Percent inhibition was calculated by normalizing measured contraction values against both positive controls (e.g. no TGF-β1) and negative controls (TGF-β1 and vehicle only).

### Pharmacological validation of My-FLECS with PIKfyve inhibitors

PIKfyve inhibitors Apilimod and YM-201636 (Sigma) were dissolved at 10mM in DMSO and arranged into a 10-step 3-fold serial dilution columnwise-wise in 3 rows each on a 384-well drug source plate. FLECSplates were pre-filled with 25 µL of serum-free DMEM, and then 100 nL of the PIKfyve inhibitor solutions were transferred from their source plate to the corresponding wells on the FLECSplate using a Biomek FXP pin tool. The FLECSplates were shaken at 100 RPM for 15 minutes to ensure mixing. HLF cells were then seeded as previously described, with TGF-β1 already mixed into the cell suspension, into columns 2-23, while column was seeded with cells not receiving TGF-β1. Plates were imaged after 24hrs as previously described. Percent inhibition was calculated by normalizing measured contraction values against both positive controls (e.g. no TGF-β1) and negative controls (TGF-β1 and vehicle only).

### Compounds

Our pilot screen was performed against the library of pharmacologically-active compounds (LOPAC1280) comprising a total of 4 384-well drug plates. Our high-throughput screen was performed against a portion of the Enamine Hit Locator Library 2021 100K set (HLL-100 2021) comprising 127 384-well drug plates. The 384-well library plates containing 10mM stock solutions were purchased frozen and remained stored at −80C until their use in the screen. Primary HTS hits were repurchased as powder from Enamine and dissolved at 10mM within 2 days of retesting.

### Primary HTS

For all screening runs, HLF cryopreserved at passage 3 were thawed into a T75 flask 7 days before the experiment. Screening runs comprised of 2 days and were executed with an integrated automation platform comprising of a Multidrop for plate-wise reagent and cell addition, a Biomek FXP for pinning drug solution, a Cytomat automated incubator and robotic arm for transporting plates between incubation and imaging stations, and a Molecular Devices ImageXpress microscope for high-throughput automated imaging.

On day 1, FLECSplates were filled with media, equilibrated to RT, given drug from the library plates (250nL of 1mM drug solution yielding a 3.5µM final concentration), then shaken on a plate shaker for 15 minutes. Next, HLF were dissociated and seeded into drug-bearing FLECSplates as previously described with TGF-β1 in plate columns 2-23 and without TGF-β1 in columns 1 and 24. On day 2, following 24 hrs of incubation, cells were given Hoechst 33342 live nuclear stain as previously described and the plates were imaged with a 4x objective at one position per well capturing both the micropatterns (TRITC channel) and stained nuclei (DAPI channel). Images were subsequently downloaded in .TIFF format and analyzed using Forcyte’s proprietary computer vision to derive quantitative contractility data.

### HTS hit criteria

Compounds were selected as hits for retesting from the primary screen using an “inhibition” cutoff off at least 75% but no more than 200% with respect to native HLF contraction. Hits were required to have a z-score on cell count greater than −1.5.

### Initial hit confirmation and triage

After completing the full 40,640 compound screen, the identified hits were cherry-picked from their original drug plates into a new confirmation drug source plate using the Biomek FXP and retested in triplicate at both 3.5µM and 0.75µM in HLF cells using the same protocol as was used for primary screening. After imaging of these experimental FLECSplates was complete, CellTiter-Glo® 2.0 Cell Viability Assay solution (Promega) was added directly to the FLECSplates for 30 minutes, per the manufacturer’s protocol, and a luminescence plate reader was used to provide a readout of cell ATP activity after 24 hrs of exposure to drug.

Cherry-picked hits were also tested in triplicate at 3.5µM and 0.75 µM against human tracheal smooth muscle (HTSM) tonic contraction in the FLECS assay. This was done using the same protocol as was used to screen compounds in HLF, but without the addition of TGF-β1 and in Ham’s F12 medium supplemented with 10% FBS rather than in DMEM.

### Hit retest from powder in dose-response

Hits deemed selective to HLF over HTSM, non-toxic, and free of undesirable structural motifs were resupplied as fresh powder from Enamine. For subsequent dose-response experiments, purchased hit compounds were dissolved in DMSO to 10mM stock concentrations, and arranged in 10-step, 3-fold serial dilutions column-wise in three rows each on new drug source plates. Testing of hits in dose-response format on HLF was performed following the same protocol as used for screening.

### Hit counter-screens in smooth muscle cells

To further phenotypically profile the hits, dose-response experiments were conducted on both HTSM tonic contraction and HTSM activation with TGF-β1. HTSM tonic contraction experiments followed a protocol similar to the HLF experiments, but without the addition of TGF-β1 and using Ham’s F12 medium instead of DMEM. Inhibition of HTSM tonic contraction was calculated by normalizing the measured values against negative controls. HTSM experiments involving activation with TGF-β1 were performed in Ham’s F12 medium with the addition of TGF-β1, following the same general procedures as the HLF experiments. Inhibition of HTSM activation with TGF-β1 was calculated by normalizing the measured values against both positive and negative controls.

### Cell viability and proliferation counter-screens

HLF and HTSM cells were seeded into black plastic-bottom 384-well plates and treated with hits in dose-response format for 72 hours. Cell toxicity was assessed using the LIVE/DEAD Viability/Cytotoxicity Kit (Thermo Fisher Scientific) according to the manufacturer’s protocol. The kit employs fluorescent dyes, calcein AM and ethidium homodimer-1 (EthD-1), to label live and dead cells, respectively. After staining, cells were imaged using a fluorescence microscope, and the percentage of live and dead cells was quantified using image analysis software. Immediately following the completion of the imaging of stained cells, CellTiter-Glo® 2.0 Cell Viability Assay solution (Promega) was added to 384-well plates, per the manufacturer’s protocol, and incubated for 30 minutes. A luminescence plate reader was then used to measure ATP activity corresponding to viable cell count in each well. Also taking into account the live/dead calculation, inhibition of proliferation conferred by drug was calculated as the percentage of the ATP activity signal normalized to negative control and subtracted from 100%.

### Smad2/3 nuclear trans-location counter-screen

Primary human lung fibroblasts (HLFs) were seeded in 96-well tissue culture plates pre-coated with a 0.2% gelatin solution. After 48h, culture media was replaced with serum-free DMEM media containing 5uM hit compounds. Cells were pretreated with compounds for 30min at 37℃ before TGF-β1 was added to a final concentration of 2.5ng/ml. Two hours after TGF-β1 addition, cells were processed for Smad2/3 staining. Briefly, cells were washed with PBS, fixed with 4% formaldehyde for 15min, and permeabilized with 0.2% Triton for 10min. After blocking with 3% BSA for 1hr, cells were incubated with rabbit anti-Smad2/3 antibody (Cell Signaling Technology, Cat #8685, diluted 1000-fold in 1% BSA) for 1hr, washed, and stained with an Alexa 594-conjugated Goat anti-rabbit secondary antibody (Invitrogen). Nuclei were counterstained with Hoechst. Cells were imaged on an ImageXpress Confocal microscope (Molecular Devices) using a 20x objective. The cytoplasmic/nuclear intensity ratio was quantified using CellProfiler^20^. Each hit was measured in triplicate.

### Precision-cut lung slice assay

Frozen human precision cut lung slices (hPCLS) from healthy donors were purchased from AnaBios. The slices were thawed, punched into 3mm discs using a biopsy puncher and cultured in DMEM/F12 media (supplemented with 0.1% FBS and antibiotics/antimycotics) in 24-well tissue culture plates. 24hrs after thawing, lung slices were pretreated with hit compounds for 1hr before adding TGF-β1 at a final concentration of 5ng/ml. Each condition was run in quadruplicate. 48hrs after treatment, slices were homogenized in RLT lysis buffer on a Bead Mill 4 homogenizer using ceramic beads, and RNA was extracted using the RNeasy Mini kit (Qiagen). Reverse transcription was performed using the High-Capacity cDNA Reverse Transcription Kit (Thermo Fisher). 4-plex qPCR for ACTB, ACTA2, Col1 and FN1 was optimized and performed using the TaqMan™ Multiplex Master Mix (Thermo Fisher) per manufacturer’s User Guide. Primer and probe sequences can be found in Supplementary Materials & Methods. Quantitative PCR was performed on a ViiA 7 Real-Time PCR System. The threshold cycle (Ct) values of ACTA2, Col1 and FN1 were normalized to the house-keeping gene ACTB and the relative expression levels were calculated using the 2ΔCt method with the average of “No TGF-β1” control samples set as 1.

### Data plotting

Prism software (Graphpad) was used to plot and curve-fit normalized data to derive IC50 values.

## Results

### FLECS Technology as a foundational platform for developing disease-specific contractility assays

To achieve HTS of contractile cell force, it is necessary to have a scalable and automated assay, where human intervention is minimal and can be replaced with automation workflows and algorithms. This requirement applies to every step of the screening process, from set-up to data output. To address this challenge, we previously developed a scalable cellular force cytometer called FLECS Technology^14^.

The original FLECS assay adopts a standard well-plate format but replaces the traditionally rigid surface with a highly compliant and elastic thin film. This film is provided with arrays of covalently micropatterned fluorescently tagged adhesive ECM proteins (Fig. 2a). Non-patterned surfaces within the wells are chemically blocked to limit cellular adhesion solely to the micropatterns. Single cells seeded from suspension self-assemble and adhere over these micropatterns, applying inward contractile forces that cause significant changes in micropattern size. Even minor size changes in the micropatterns (∼1 µm) are visible under low magnification microscopy (e.g., 4X) and can be accurately quantified using computer vision algorithms (Fig. 2b). The precision of the micropatterns’ shape, size, and orientation minimizes environmental variation across individual cells, ensuring consistent cell footprint and orientation. Micropatterning also immobilizes cells without affecting their contractility, facilitating the alignment of images taken at different time points and enabling the capture of time-dependent contractile changes in each cell. A live nuclear dye is added at the final time point to aid in the detection and counting of cells across every identified micropattern (Fig. 2c). Contractility quantification is performed on a per-cell basis and normalized to the original micropattern dimensions sourced from unoccupied reference micropatterns in the same image. This process provides a standardized scale for normalizing and comparing measurements across large screens. The characteristics of the FLECS system enable straightforward automation of cell seeding, plate handling, imaging at only one wavelength during live recordings, and data analysis.

**Fig. 2:**
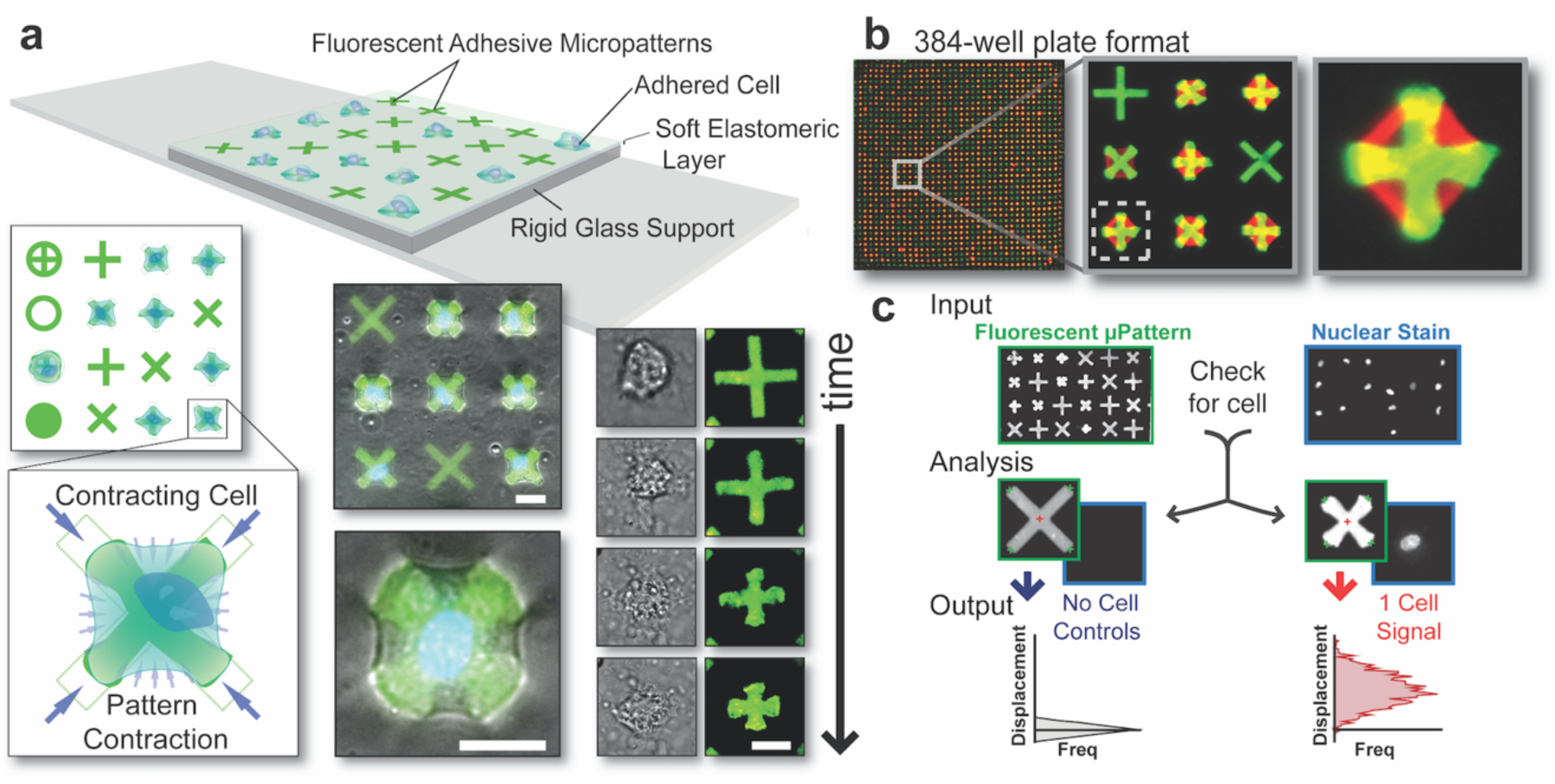
a) Schematic representation of the FLECS technology: Single-cells adhered to adhesive micropatterns embedded within an elastomeric film exert mechanical force onto the micropatterns resulting in their displacements. Top view demonstrates various pattern shapes, while a magnified image illustrates a contracting cell inwardly displacing its terminals. Overlay of fluorescent patterns and phase contrast images captures contracting cells, with time-lapsed images showcasing the cell’s contraction over the underlying micropattern. Scale bars represent 25 μm. b) Implementation of FLECS Technology in a 384 well-plate format. c) Workflow for image analysis: Input includes aligned image sets of the micropatterns (set 1) and stained cell nuclei (set 2). Analysis involves (i) identification and measurement of all micropatterns in image set 1, (ii) cross-referencing the positions of each micropattern in image set 2, and (iii) determining the presence of 0, 1, or >2 nuclei (cells). Output consists of mean center-to-terminal displacements of micropatterns containing a single nucleus (one cell), compared to the median measurement of non-displaced patterns with zero nuclei. The resulting differences are presented as a horizontal histogram. Figure adapted with permission from Pushkarsky *et al*^14^.

Here, we transferred the platform to a 384-well multi-well format meeting the requirements for industrial HTS and allowing the screening of contractile cell force of up to 100,000 compounds per month. At the 384-well level, a 4x objective will capture an entire well within a single field-of-view, while still achieving the required image resolution for accurate analysis due to the precise geometries of the micropatterns. This allows for imaging of an entire 384-well plate to complete in under 10 minutes using a high-content imaging microscope, while only generating a data capacity of only 18 MB per well. Using primary human lung fibroblasts (HLF) as our model fibroblast cell type, we developed a FLECS-based myofibroblast contraction assay (My-FLECS) that can rapidly identify and quantify inhibitors of myofibroblast contractility following activation with TGF-β1.

### Initial development and validation of a functional contractility assay reporting activation of primary HLF using FLECS Technology

We adapted the core FLECS platform specifically for myofibroblasts, creating My-FLECS, through a series of optimization studies examining critical system parameters such as micropattern size, film stiffness, micropattern protein, and culture media conditions. The results demonstrated that 70µm ‘X’ shaped micropatterns, composed of type IV collagen on approximately 8 kPa stiffness films, effectively promote both fibroblast adhesion and contractile response. In order to establish activated (e.g. disease-driving) and non-activated control baselines for the My-FLECS, cell contractility was compared between HLFs cultured with and without 2.5 ng/mL TGF-β1 for 24 h (Fig 3a). A 10-step 2-fold titration of TGF-β1 confirmed 2.5 ng/mL as the optimal dose for activating myofibroblasts in the My-FLECS assay (Fig. 3b). Relative to the low baseline substantial tonic contractile forces of untreated HLFs, TGF-β1-*activated* HLFs generated between 50% to >80% larger micropattern displacements on average (depending on cell passage number). Analysis of the single-cell distributions for each group showed that the relatively wide normal distribution of untreated HLF contraction measurements transitions to a much narrower distribution in activated HLF. Thus, in addition to producing HLF with significantly higher contractile force, TGF-β1 treatment resulted in greater cell contraction homogeneity. Both results are consistent with prior reports that fibroblasts are heterogeneous with respect to their spontaneous activation *in vitro*^21^, as reflected by the wide distribution for untreated HLF, and that TGF-β1 treated fibroblasts become highly contractile^22^. During the optimization of My-FLECS, we observed that TGF-β1 responsiveness was maximized in fibroblast growth factor (FGF)-free conditions. Unique to HLF, compared to other cell types tested with FLECS, was a heightened sensitivity to pH variations in the medium, necessitating the need for precise pH buffering.

**Fig. 3:**
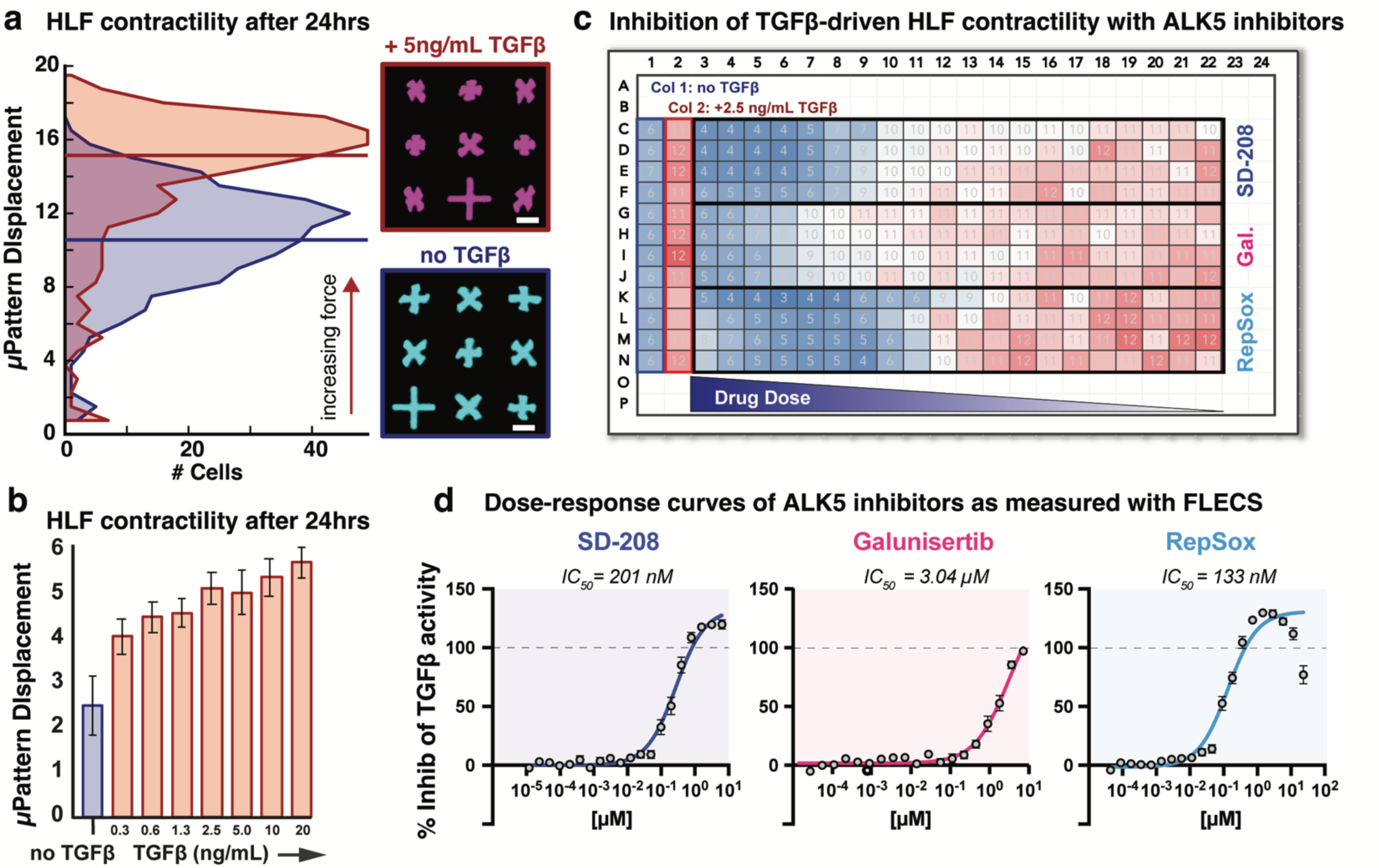
Development and validation of a myofibroblast contractility assay based on FLECS Technology. A) Representative distributions of single-cell contractility in HLF grown within wells absent TGF-β1 (blue) or with 2.5 ng/mL TGF-β1 treatment (red). Treated cells demonstrate significantly increased contraction after 24 hours. Corresponding blown-up images of the two populations are displayed. B) Concentration-response analysis of TGF-β1 effects on HLF contractility reveals a dose-dependent activation. C) Heatmap representation of an experiment assessing the dose-response of ALK5 inhibitors on HLF activation using the established assay. Column 1 represents the absence of TGF-β1. Column 2 represents 2.5 ng/mL TGF-β1 with DMSO only. A twenty-step, 2-fold concentration response array is presented from column 3 (15 µM) to column 22. D) Concentration response curves of the heatmap data shown in (C). As shown, four technical replicates were used for each dose of each drug. Each replicate comprises a 384-well of approximately 400 HLF single-cells.

To validate the ability of our assay to identify inhibitors of HLF activation, we next performed a dose-response experiment characterizing the effects of three known inhibitors of the TGF-β1 receptor I (RI) ALK-5 on HLF contractility following 24 h treatment with TGF-β1. Following a 20-step 3-fold dilution series, the three ALK-5 inhibitors (SD-208, Galunisertib, and RepSox) were organized into a single drug source 384-well-plate (Fig. 3C) for simultaneous assessment. As expected from TGF-β1-R1 inhibitors, exposure of HLF to all three ALK-5 inhibitors for 30 min prior to adding TGF-β1 inhibited activation of HLF into highly contractile cells.

### Further assay validation with inhibition of recently identified intracellular targets regulating fibroblast activation

Although exceptionally effective in preventing TGF-β1-driven myofibroblast activation, the direct blockade of TGF-β1 or of its receptor is not a viable clinical approach to therapeutically treat fibrosis due to the systemic importance of TGF-β1 signaling^23^. Thus, we next sought to further validate our assay’s ability to detect inhibition activity occurring in pathways downstream of TGF-β1-TGF-β1-RI binding. We selected two inhibitors of the kinase PYKfyve (apilimod and YM-201636) which were recently shown to modulate fibrotic activation *in vitro* and *in vivo*^24^.

We utilized the FLECS platform to evaluate the effects of these PIKfyve inhibitors on both HLF activation with TGF-β1 and, in parallel, on the basal contractile tone of primary human tracheal smooth muscle (HTSM) cells. HTMS cells were chosen since they also reside within the human respiratory system and are phenotypically similar two myofibroblasts. Successful treatments of fibrosis ideally target myofibroblasts without affecting other contractile cells in the target organ like HTMS (lung), vascular smooth muscle (all organs), myocytes (muscle), or cardiomyocytes (heart). PIKfyve inhibition prevented TGF-β1-driven activation of HLF but did not antagonize basal contraction of tracheal smooth muscle (Fig. 4). Hence, the My-FLECS assay is not only able to identify inhibitors of HLF activation downstream of TGF-β1R1 binding but can also differentiate between general effects on cell contractile mechanisms and myofibroblast-specific contraction. With these successful demonstrations, the assay met the criteria for use in anti-fibrotic drug discovery and was ready for scale-up.

**Fig. 4:**
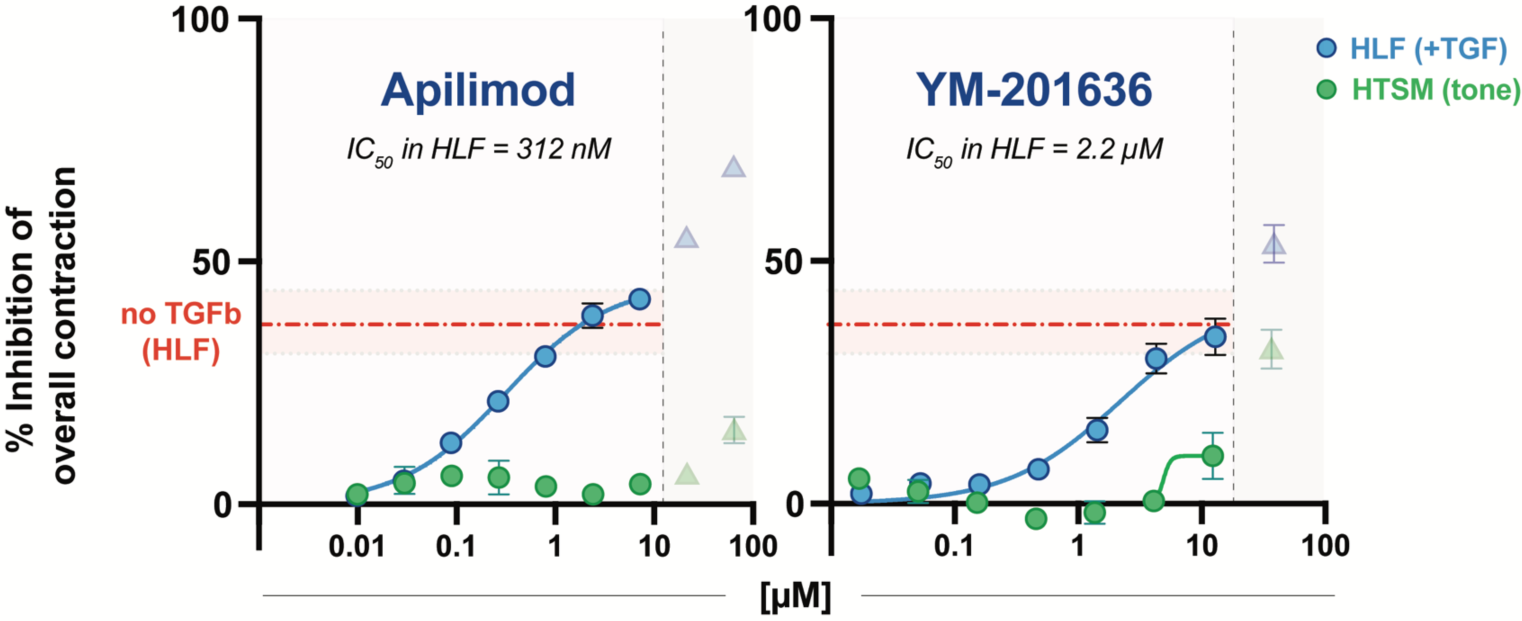
Dose-response analysis of the effects of PIKfyve inhibitors apilimod and YM-201636 on TGF-β1-driven contractility in HLF (blue) and on tonic contractility in HTSM (green). Y-axis represents normalized inhibition of overall contraction such that 100% corresponds to a complete absence of contractile force. The red dashed lines designate the contractile level of native HLF in terms of percent reduction from TGF-β1-treated levels. Both inhibitors were able to inhibit TGF-β1 driven HLF contraction in a dose-dependent manner without any measurable effect on HTSM tone across a range of non-toxic doses. However, at doses exceeding 10µM, apparent toxicity, characterized by cell de-adhesion from micropatterns, was observed in both cell types. Although these data are presented, they were not included in curve fitting and analysis. For HLF, each data point shown comprises 9 technical replicates pooled from 3 different assay plates. For HTSM, each data point shown comprises 3 technical replicates. Each replicate comprises a 384-well of approximately 400 single-cells.

### Set-up of a HTS protocol to identify inhibitors of contractility in activated fibroblasts

To achieve true HTS campaigns with the My-FLECS platform, we next developed an automated workflow to screen up to 15,000 compounds per day, equating to approximately 48 384-well plates. Given the relatively low number of reagent additions and wash steps required by our assay, as compared to traditional high-content screening workflows where numerous solution exchange steps are required for fixing, blocking and multi-channel staining of cells^25^, as well as the native well-plate format of the assay, our automation strategy was straightforward (Fig. 5). FLECS 384-wellplates were manufactured in batches by Forcyte Biotechnologies and stored with sterile PBS at 4°C covered with aluminum seals until their use. On the first day of a screening experiment, automation systems were utilized to pre-fill each plate with cell growth medium, add compounds to the plates using a pin tool, and seed HLF cells in medium containing TGF-β1. Following an overnight incubation in the presence of the drug and TGF-β1, the automation system was used to deliver a live nuclear stain and perform high-throughput imaging of all plates.

**Fig. 5:**
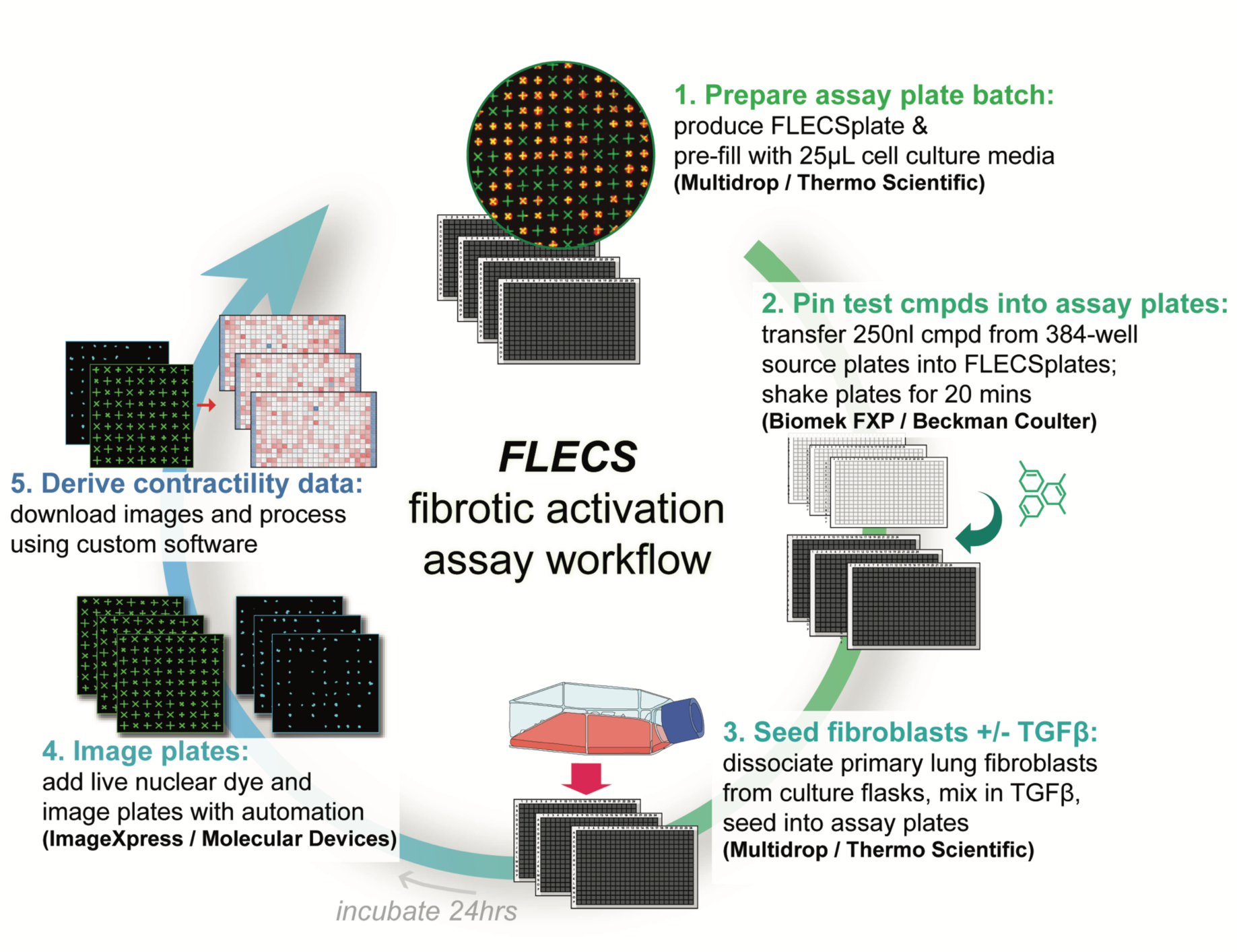
Workflow and equipment used to automate the developed assay for HTS campaigns.

### Pilot screen of a library of pharmacologically-active compounds (LOPAC) as validation for HTS

After establishing the necessary automated workflow, we evaluated the performance, variability, robustness, and reproducibility of the assay by screening the complete LOPAC1280 library at a single concentration (3.5 µM) in duplicate across 8 384-well plates in a single batch (Fig. 5). Columns 1 and 24 represented non-activated native HLF as a positive control, while columns 2 and 23 were TGF-β1-treated and received vehicle only, serving as negative controls for uninhibited myofibroblast activation. The robust Z-prime values ranged from 0.4 to 0.7 across the 8 plates. We defined ‘hits’ as compounds limiting contraction in TGF-β1-treated HLF by at least 75% but no more than 200% with respect to native HLF contraction. Additionally, to eliminate potentially toxic compounds, hits were required to have a z-score on cell count greater than −1.5. Fig.6 shows the reproducibility of the assay and consistency in hits when run in an automated batch and thus, indicates that the assay is suitable for single compound concentration screening.

**Fig. 6:**
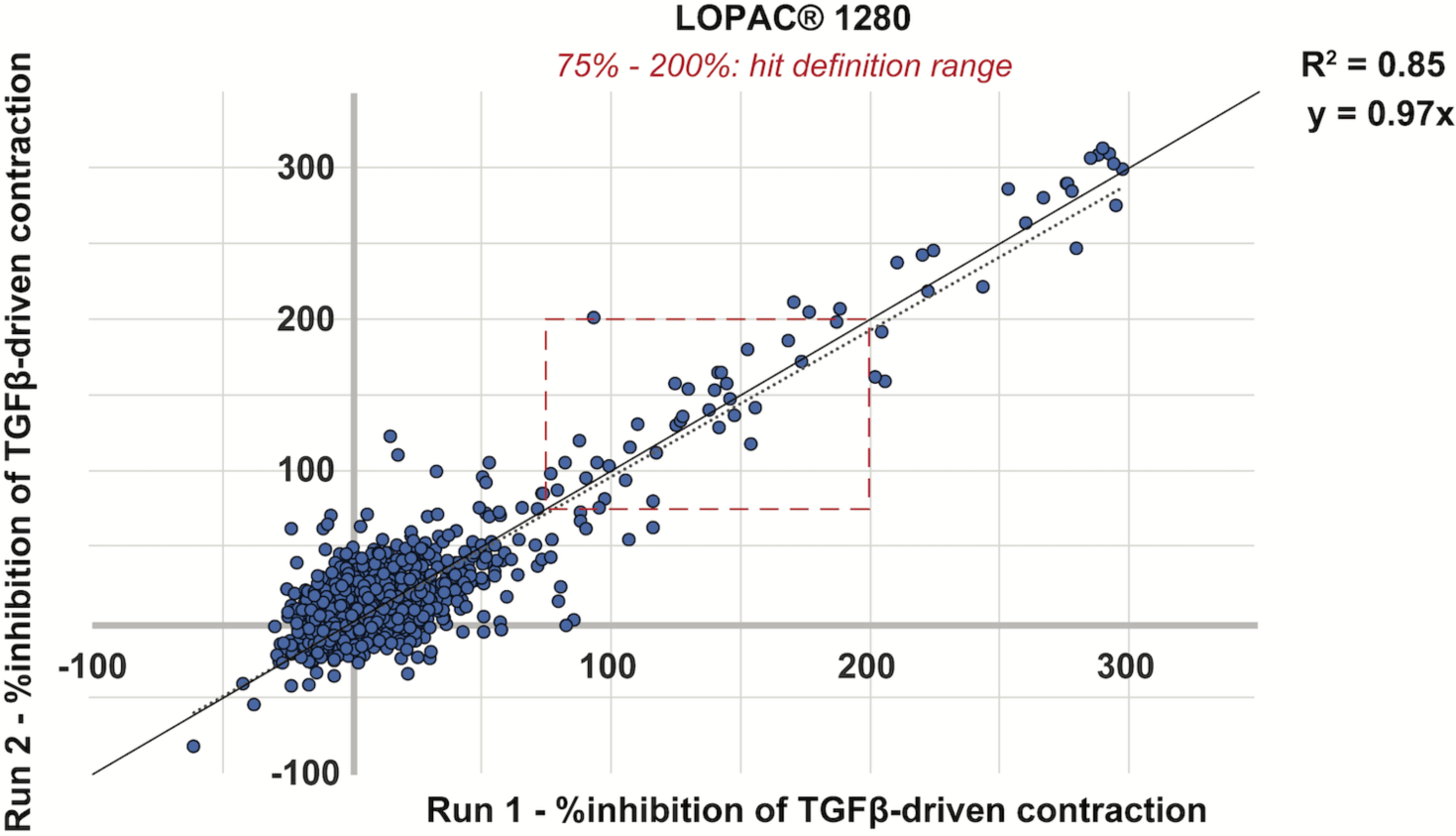
Assay reproducibility. Activities of the compounds found in the LOPAC1280 library tested twice within the same automation batch are plotted. Solid lines represent a 1:1 correlation of percent activity, and the dashed line represents line of best fit. Here, 100% inhibition of myofibroblast activation is defined as a reduction in contraction of TGF-β1-treated cells to levels equivalent to non-treated cells, as non-treated cells exhibit a baseline mechanical force upon adhering to the ECM. Compounds that would be considered hits appear within the dashed-line box representing activity levels of 75%-200%. Satisfactory reproducibility is observed.

### Rapid primary HTS with >40,000 compound library in primary HLF and counter screening

Satisfied with the reproducibility and overall performance from the LOPAC1280 screen, we next implemented the same automation strategy to execute out a HTS of a larger commercial sub-library. The goal was to identify additional compounds that could inhibit TGF-β1-induced HLF activation, thus providing potential starting points for developing anti-fibrotic therapeutics. In total, we screened 40,640 compounds distributed across 5 experimental batches, each containing between 10 and 45 assay plates.

To ensure the assay performance was robust, the batches were incrementally scaled up, starting with 10 plates per batch and scaling up to 45 plates in the fourth batch. To accommodate the larger batch size, two automation stations were used in parallel to image the assay plates. For the fifth and final batch, a single automation station was used to screen 40 plates. Here, the longer exposure of cells to the nuclear stain in plates towards the end of the batch resulted in a larger variation in robust Z-prime values compared to the first four batches (Fig. 7a). Nonetheless, the performance of the assay was still satisfactory for a cell-based functional-phenotypic screen. The performance metrics for each batch also highlight the batch size (Fig. 7b). These hits represented potential starting points for further development as anti-fibrotic therapeutics.

**Fig. 7:**
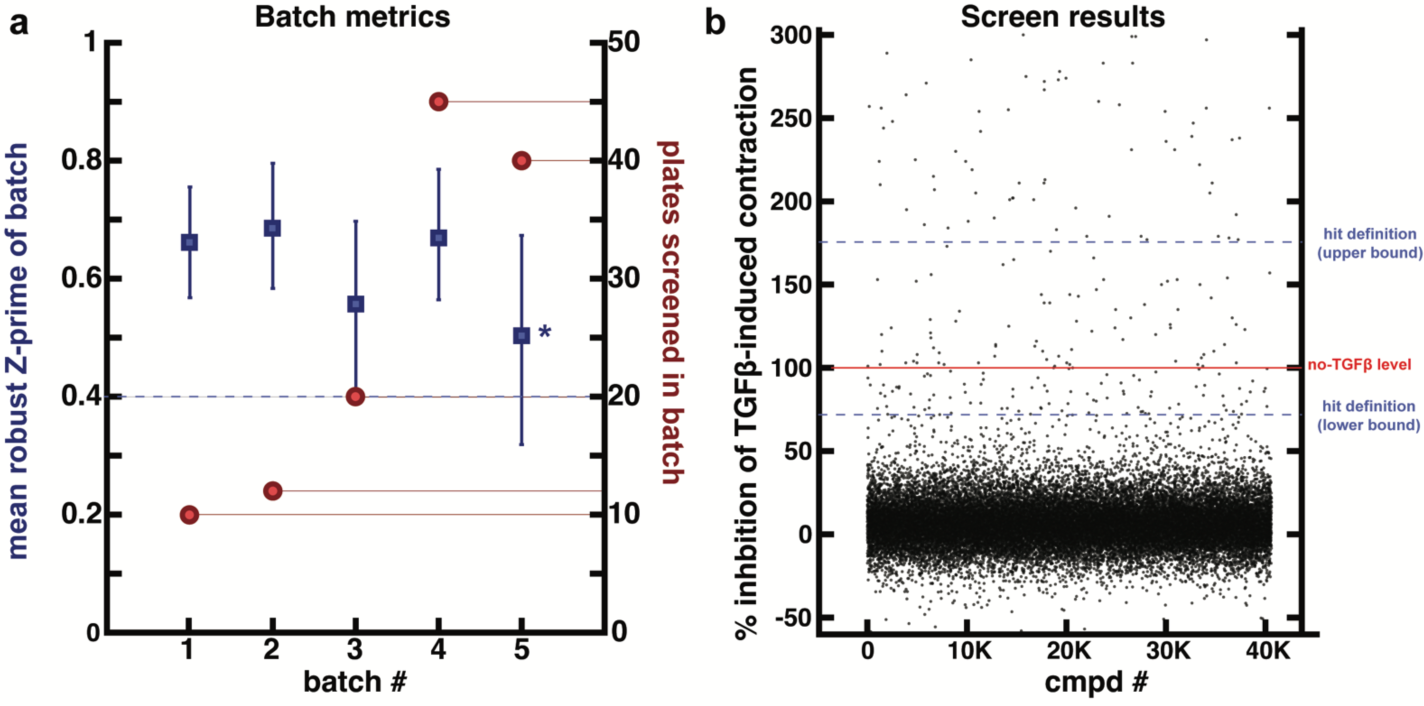
HTS campaign metrics. A) Batch metrics for the screening of 40.6k compounds. The screening process was completed in five batches, with batch sizes progressively increasing (indicated on the right axis). Note that Batch 4 was imaged across two automation stages in parallel. Batch 5, imaged at a single station, took longer to complete overall and resulted in longer exposure of cells seeded on plates imaged at later times to nuclear stain. Consequently, some plates exhibited a lower than typical robust Z-prime value (indicated on the left axis). B) Data scatter plot showcasing the results for all compounds screened. The blue dashed lines indicate the boundaries defining hits in terms of the percentage inhibition of TGF-β1-driven contraction in HFL. The red solid line represents the contractile level of native HLF, defined as 100% inhibition.

Overall, 166 initial hits were identified in the primary screen corresponding to a hit rate of 0.4%. We then used automation to cherry-pick hits samples from the original compound library into a new ‘hit’ source plate for confirmation screening which was conducted in triplicate at both the original screening dose of 3.5 µM and a reduced dose of 0.75 µM. Since the primary screen only provided a phenotypic profile of the hits in the target HLF cell type, we also used the FLECS assay to counter-screen the cherry-picked hits against contractile tone in HTSM cells. The hits that were confirmed at the original dose, active at the lower retest dose, and inactive in HTSM cells were further evaluated for known structural liabilities such as reactive groups, PAINS structures, and cytotoxic motifs. Through this triage process, we narrowed our final list down to 42 compounds, which were repurchased as powder and tested in a 10-step, 3-fold concentration-response assay ranging from 1 nM to 35 µM.

Concentration-response testing of resupplied hits was conducted in both the original HLF assay and in a panel of phenotypic counter-screens reporting on cell viability (live/dead), proliferation (ATP assay), and off-target effects on HTSM tonic contraction over a broader dose-range (FLECS assay). Moreover, given that cultured HTSM respond to TGF-β1 like HLF by becoming significantly more contractile^26^, we also counter-screened the hits resupplied as powder in a modified HTSM contractility assay where TGF-β1 was used as a pro-contractile trigger. The goal of this counter-screen was to determine if the primary hits, which were originally found in HLF, inhibited broad TGF-β1 signaling pathways shared by multiple cell types or if they were selective to pathways within HLF. Selectivity to pathways within HLF was considered more attractive and novel, given the safety issues facing broad-spectrum ALK5 inhibitors in therapeutic development. The entire process of screening 40,460 compounds, confirming hits, and performing deep phenotypic characterization across multiple assays and cell types on confirmed hits was completed in a mere 12 weeks.

### My-FLECS identifies novel anti-fibrotic hit molecules

After eliminating compounds that exhibited adverse effects on cell viability antagonized HTSM contractility, or had low potency (<1µM), the remaining hit set comprising 19 molecules displayed a promising pharmacological profile that supported further development. This profile is distinguished by a considerable reduction in TGF-β1-driven contractility in HLF cells but not in HTSM cells, a marked inhibition in proliferation of HLF cells (without affecting viability, Supplemental Data) but not in HTSM cells, and a minimal impact on baseline contractile tone of HTSM cells. Activity profiles are shown for four most potent hits in Fig. 9A.

**Fig. 8:**
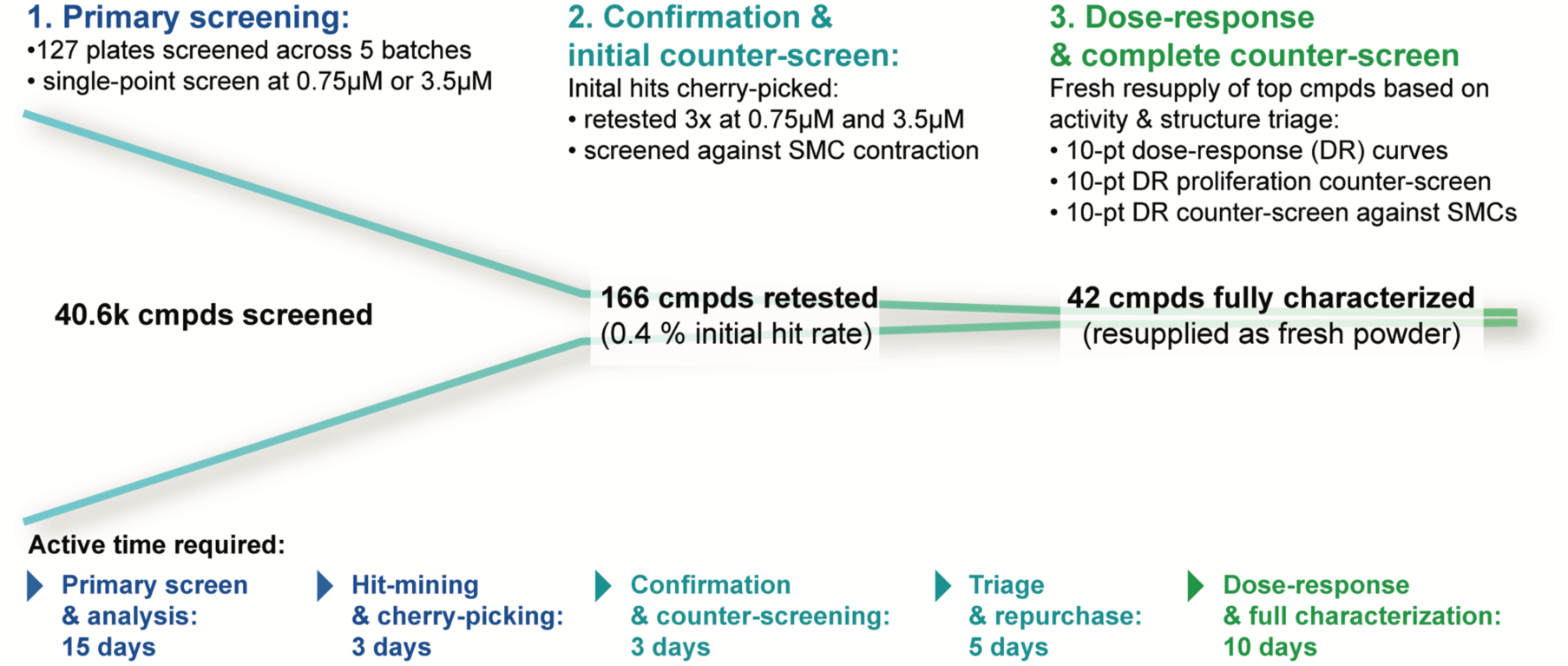
Compound screening progression. Our strategy for identifying and prioritizing hits for repurchase following a 40.6k screen is shown. We screened 127 384-well library plates (corresponding to 40.6k compounds) at a single dose of 3.5µM across 5 batches as previously described over a 5-week period inclusive of completing automated data analysis on a cloud server. We next cherry-picked and retested 166 initial hits in both HLF and HTSM cells 3 times each at multiple doses. Hits exhibiting consistent and selective activity without obvious structural liabilities were repurchased as fresh powder. Upon receipt, confirmed hits were re-tested in a dose range in the core assay and in several counter-screens. The overall process, including waiting periods, took a total of 15 weeks and ultimately yield 19 potent hits with extremely attractive phenotypic profiles.

**Fig. 9:**
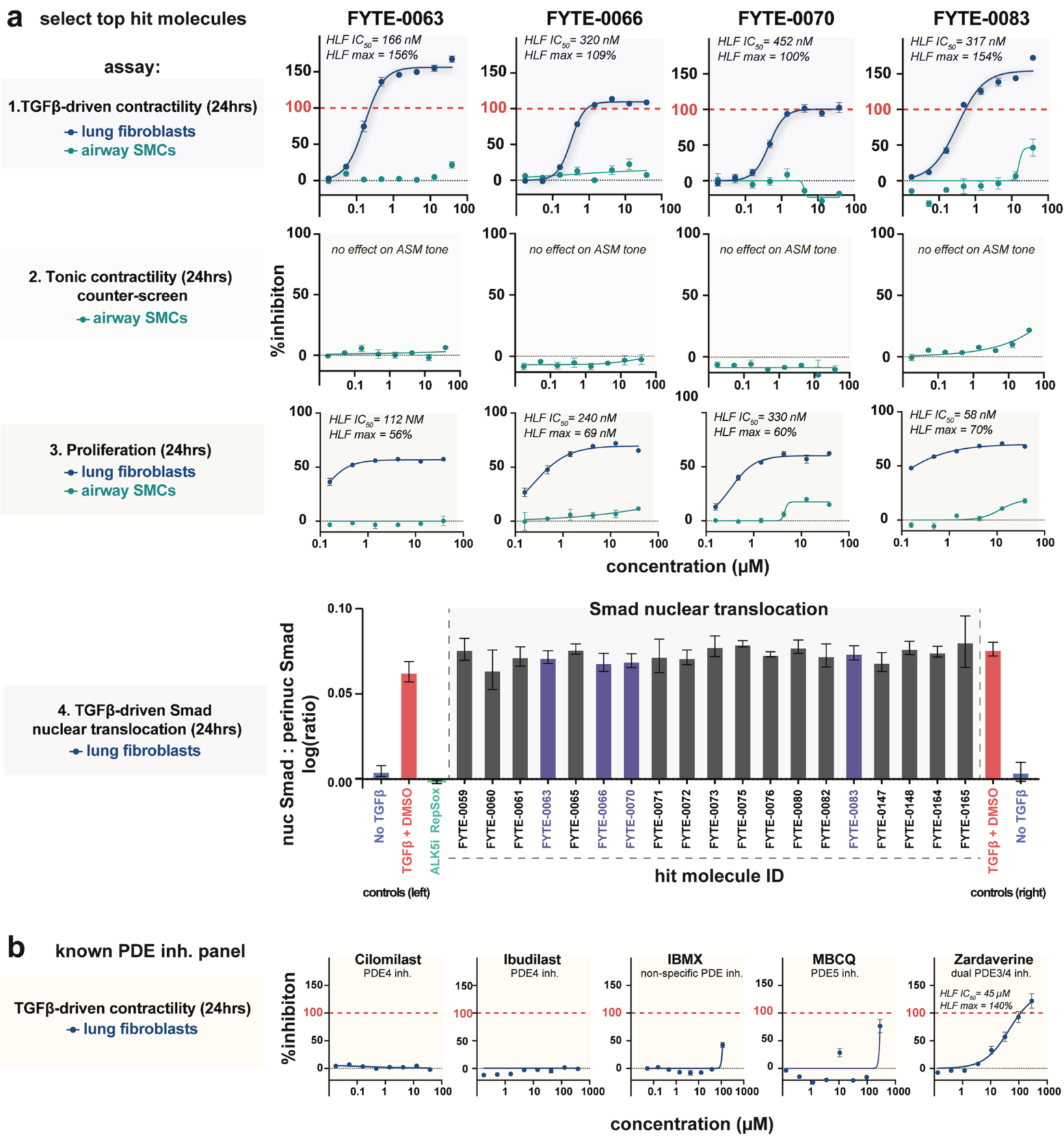
Representative anti-fibrotic hits from our screening campaign. (A) Four of the top 19 hits are showcased, all sharing a phenotypic profile characterized by potent inhibition of TGF-β1-induced contractility in human lung fibroblasts (HLF) without similar effects in human tracheal smooth muscle cell (HTSM) (row 1), no measurable influence on HTSM contractile tone (row 2), strong suppression of HLF proliferation without impacting viability or affecting HTSM (row 3), and no attenuation of Smad nuclear translocation (row 4). Data from the Smad nuclear translocation assay is presented for all compounds sharing this phenotypic profile. The representative hits displayed in rows 1-3 are highlighted in violet. (B) Panel of PDE inhibitors tested in the core HLF contractility assay based on structural similarities to some of our hits. The results suggest that despite apparent structural similarities, PDE inhibition does not appear to be the mechanism by which our hits have effect, suggesting potentially novel mechanism are involved. For HLF contraction, each data point shown comprises 9 technical replicates pooled from 3 different assay plates. For all HTSM contraction and all proliferation data, each data point comprises 3 technical replicates. Each replicated comprises a 384-well with approximately 400 single-cells. Smad translocation data comprises 3 technical replicates each consisting of a distinct well with >100 cells evaluated from in the field-of-view.

To address potential mechanisms of action of our advanced hits, we probed whether they acted upstream or downstream of the TGF-β1-mediated nuclear translocation of Smad2/3 by employing fluorescent staining and assessing the Smad signal ratio in the nuclear and perinuclear regions 24 h following TGF-β1 exposure. Inhibition of Smad nuclear translocation would imply interference with TGF-β1 receptor engagement or processes immediately following it while actions independent of Smad signaling would indicate interference with non-canonical pathways. In contrast to our control ALK5 inhibitor (RepSox), none of our hits affected TGF-β1-stimulated Smad nuclear accumulation (Fig 9a-4). Thus, our compounds act downstream of the TGF-β1-RI or through non-canonical pathways with potential selectivity towards HLF-to-myofibroblast activation.

Next, we assessed chemical structure resemblances of our advanced hits with that of annotated compounds with established activities by reviewing the CHEMBL database, scientific literature, and patents. Our analysis served to evaluate the novelty of our findings and to generate potential target hypotheses based on the identified structural similarities. Most of our hits contained a prevalent structural motif common to nitrogen-containing heterocycles found in diverse phosphodiesterase (PDE) inhibitors. To test whether PDE inhibition is a potential mechanism of preventing fibroblast activation, we performed a concentration response analysis of a panel of tool inhibitors of PDE4, PDE5, a pan-PDE inhibitor, and a dual inhibitor of PDE3/4 (Fig. 9a). Among the tested PDE inhibitors, only the dual-selective PDE3/4 inhibitor Zardaverine exhibited activity in our My-FLECVS assay, reaching and exceeding 100% inhibition above 100 µM. The calculated IC50 for this response was approximately 45 µM, which is nearly three orders of magnitude higher than the IC50 values reported for its known PDE targets (170 nM and 580 nM to PDE3 and PDE4, respectively). Zardaverine also did not exhibit an obvious plateau in concentration-response curve even at high doses above 100 µM. Further considering that two other PDE4 inhibitors failed to affect TGF-β1-induced HDF contraction, PDE3/4 inhibition is unlikely the mechanism through which Zardaverine inhibiting fibroblast activation and contraction. Collectively, these results suggest that our advanced hits act through a potentially novel mechanism of action, unrelated to inhibition of ALK5 binding, pSmad shuttling, PDE activity, and may therefore represent valuable new therapeutic opportunities.

### Identified hits suppress fibrotic gene upregulation in human precision-cut lung slices

Finally, we performed initial translational testing on our advanced hits by evaluating their ability to suppress TGF-β1-driven upregulation of fibrotic gene expression in human precision-cut lung slices (hPCLS) (Fig 10a). Because our best hits inhibited TGF-β1-induced rather than baseline contractile function, we hypothesized that they exert broader effects on markers associated with TGF-β1-driven myofibroblast activation. To pursue this idea, we assessed whether our compounds prevented the upregulation of the canonical myofibroblast contractile protein α-SMA (encoded by ACTA2), and the ECM protein collagen I (COL1A1), another myofibroblast product in fibrotic tissues.

**Fig. 10:**
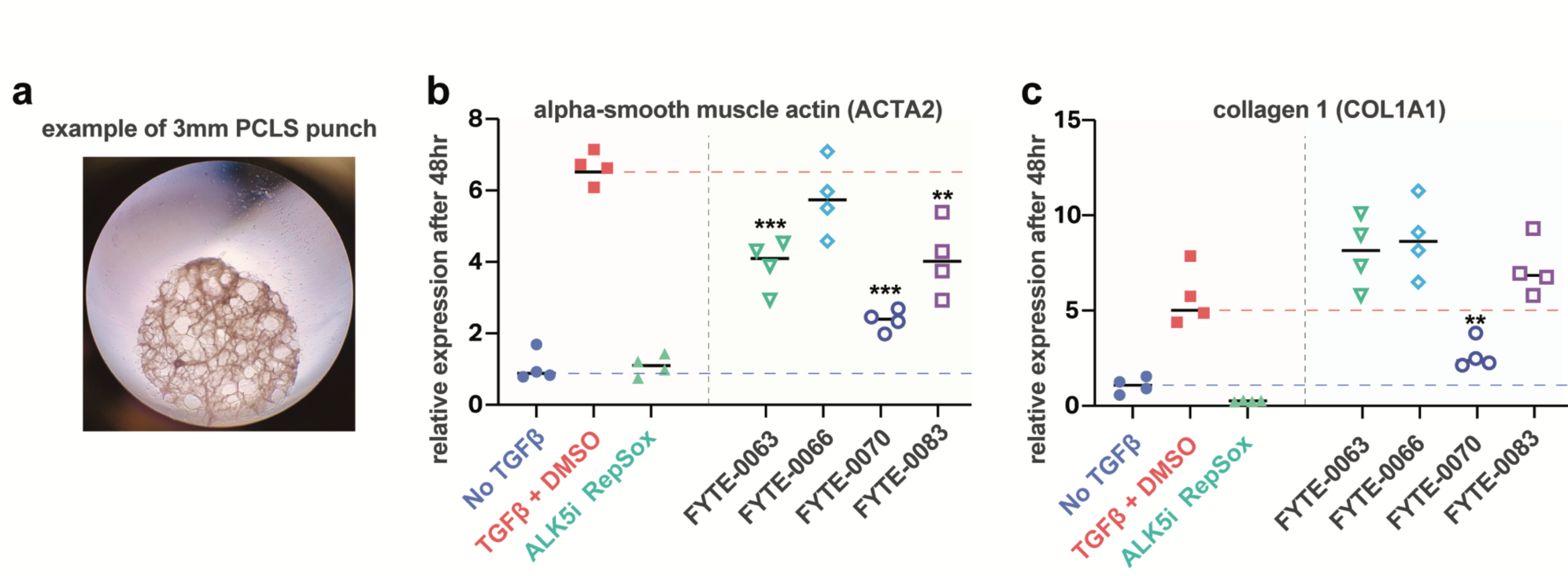
Translation studies on hits using human precision-cut lung slices (PCLS). (A) Photograph of a 3 mm punch taken from a 250 µm thick human lung slice within a well on a 96-well plate. Four punches were utilized for each test condition. (B) Relative transcript levels of ACTA2 following treatment with the listed conditions. Three out of the four hits tested in this model exhibit a significant reduction in ACTA2 upregulation compared to the controls. (C) Relative transcript levels of COL1A1 following treatment with the listed conditions. Unexpectedly, one of the tested hits, originally identified for its activity in suppressing TGF-β1-driven HLF contractility, was also found to suppress the upregulation of COLA1. Each data point comprises a unique biological replicate (e.g. a different 3mm tissue punch treated in separate well on a plate). For each biological replicate, qPCR reactions were performed using four technical replicates.

We observed a substantial increase in the transcript levels of both ACTA2 and COL1A1 in hPCLS treated with TGF-β1 for 48h, compared to untreated hPCLS (Fig. 10a). This upregulation was effectively abrogated by co-incubation with the ALK5 inhibitor RepSox (Fig. 10a). Of the four tested hit compounds identified in our My-FLECS screens, three exhibited significant inhibition of ACTA2 upregulation (Fig. 10b). Moreover, one of our hits also substantially inhibited upregulation of COL1A1 expression relative to controls (Fig. 10c), despite having no prior evidence predicting this activity.

## Discussion

The high contractility of activated myofibroblasts plays a pivotal role in the advancement of fibrosis, signifying its importance when formulating therapeutic strategies. Nevertheless, HTS for anti-fibrotic compounds has traditionally prioritized cellular protein expression or proliferation, often sidelining myofibroblast contractility until later stages of hit validation. To address this gap, we have developed a HTS functional assay specifically tailored to assess cellular contractility, specifically in primary human fibroblasts. This innovation enables us to concentrate on the crucial contractile phenotype in fibrosis from the very onset of the discovery process, enhancing our ability to pinpoint functionally active compounds right from the primary screening stage, providing a more targeted and efficient approach to anti-fibrotic drug discovery.

This My-FLECS assay was built on the previously reported FLECS Platform for single-cell contractility, an approach presenting several distinctive capabilities that are essential for high-throughput implementation of this phenotypic readout, including normalization of cell behavior, deployment in a 384-wellplate format, automated image analysis, and the generation of statistically significant data throughput per test condition. Using the My-FLECS, we successfully demonstrate that TGF-β1-driven contractile function can be quantified and used as a functional biomarker of myofibroblast activation in primary HLF, with excellent dynamic range and reproducibility both intra- and inter-plates. We further demonstrated the potential of the My-FLECS to be scaled to industrial HTS of >17K wells per day (45 384-well-plates), and its ability identify promising phenotypic hits from expansive drug libraries. Through comprehensive counter-screening of these hits in other cell types and assay paradigms within the core FLECS assay, we demonstrate the ability to rapidly profile additional relevant phenotypic activities, enabling data-driven hit prioritization strategies to guide the advancement of compounds with the most promising profiles. Additionally, our follow-up experiments using precision-cut lung slices provide compelling evidence that our approach, focused on isolating activity on TGF-β1-driven contractile function in activated myofibroblasts, encompasses a wide range of potential targets. As a result, it enables the identification of compounds with broad activities across various TGF-β1-driven phenotypes, highlighting the versatility and efficacy of our methodology.

During the scale-up of our assay to >24 plates per day, we encountered bottlenecks that potentially impact plate quality and overall screening outcomes. First, we observed that HLF, compared to other cell types, displayed heightened sensitivity to minor changes in medium pH before adhesion. Inadequately buffered medium led to compromised cell adhesion to the adhesive micropatterns, resulting in incomplete spreading, reduced overall contraction, and ultimately, decreased long-term cell viability. To mitigate this sensitivity, it is crucial to prepare fresh and appropriately buffered cell medium for each screening batch and minimize exposure of the medium to the ambient environment. Even a brief delay, such as holding medium in narrow tubing of a liquid dispenser for any period longer than it takes to prime the tubing, could induce observable colorimetric changes in pH, affecting cell behavior. Therefore, strict adherence to these protocols is necessary to ensure reliable and accurate results in HTS.

Secondly, we observed a significant adverse effect on the magnitude of HLF contractility with prolonged incubation using live nuclear stain, such as Hoechst 33342. Specifically, contractility measurements taken later than approximately 6 h after stain exposure exhibited a decrease in overall magnitude compared to earlier time points under identical conditions. As the post-exposure time continued to increase, overall contraction diminished to an extent that began to comprise the separation between controls, thereby reducing Z-primes and impacting the interpretability of the assay. To mitigate this interference, screening batches should be appropriately sized or divided into sets where the addition of nuclear stain is done set-wise, preventing prolonged exposure and preserving the integrity of the assay results.

Third, we occasionally observed the occurrence of “edge-effects,” commonly seen in cell-based assays, which can manifest as non-uniform cell distributions in the outer wells or increased contraction along the outer edges compared to the interior wells. To ensure consistent and uniform cell adhesion across all wells, it is crucial to seed plates that have fully equilibrated to room temperature. Temperature disparities across the plates can induce micro-flows that drive cells towards one side of the wells, leading to non-uniformity in cell adhesion distributions. By allowing plates to reach equilibrium with the ambient temperature, we can mitigate the occurrence of these edge-effects and maintain uniform cell behavior throughout the assay. Other limitations to consider are that the assay measures contractility in an isolated system, which may not fully replicate the complex *in vivo* environment where myofibroblasts interact with other cells and the extracellular matrix. We also note that while the assay showed consistency and reliability in our hands, further validation in other laboratories would be beneficial to establish its robustness. In particular, our screens and follow-up studies were performed in well-characterized cells derived from only a single donor for each unique cell type used in the study.

In conclusion, this study introduces the first ever HTS paradigm that offers the ability to identify effectors of myofibroblast contractility from large-scale primary screening campaigns, in an efficient and scalable manner that can produce promising functionally validated starting points for drug discovery. Our hPLCS results highlight the potential drawbacks of depending solely on molecular readouts, such as the quantification of ACTA2 protein levels, which are commonly employed in HTS for anti-fibrotic compounds. As demonstrated with FYTE-0066, a lack of observable activity in these molecular readouts does not definitively equate to an absence of functional efficacy in inhibiting fibrotic processes. This discrepancy indicates that a singular focus on molecular markers may inadvertently exclude compounds that could serve as promising starting points for deeper investigation. Our results further highlight the value of incorporating a functional readout into compound screening, specifically one that gauges the multi-faceted hypercontractile phenotype triggered by myofibroblast activation. As seen with, e.g., FYTE-0070, this approach facilitates the discovery of compounds that not only exhibit vital functional activity in impeding the mechanical mechanisms underlying fibrosis, but it may also reveal broader inhibitory effects on a variety of phenotypes driven by TGF-β1, extending beyond the contractile pathways. By integrating a functional viewpoint into the screening process, it’s possible to identify compounds with diverse activity, offering significant promise as potential anti-fibrotic agents.

Ultimately, this functional assay can potentially be applied to various types of fibrosis, since myofibroblast contractility plays a significant role in many fibrotic diseases in many tissues. It can also extend to other modalities including screens for biologics as well as fibrotic genes via arrayed genome-wide knock-out screens.

## Acknowledgements and funding

The research of BH is supported by a foundation grant from the Canadian Institutes of Health Research (#375597) and support from the John Evans Leadership funds (#36050, #38861, and 38430) and innovation funds (‘Fibrosis Network, #36349’) from the Canada Foundation for Innovation (CFI) and the Ontario Research Fund (ORF). The authors also thank the Magnify Incubator at CNSI and the Nanolab at UCLA, and the Molecular Screening Shared Resource at CNSI for providing critical infrastructure to support the work.

## Author Contributions

I.P. and Y.W. conceived of the project. Y.W., E.C., R.H. and J.W. performed the experiments. E.C., Y.W., and I.P. analyzed the data. R.D. developed the automation strategy and oversaw the robotic laboratory and lead compound management. I.P., Y.W., and R.D interpreted the data. I.P., B.H. and R.D. wrote the manuscript.

