## Supplemental Data for "FLECS Technology for High-Throughput Screening of Hypercontractile Cellular Phenotypes in Fibrosis: A Function-First Approach to Anti-Fibrotic Drug Discovery"

[1]Forcyte Biotechnologies, Inc. 570 Westwood Plz Bldg 114, Los Angeles, CA 90095

[4]University of California, Los Angeles, Los Angeles, CA, United States.

### Table S1: hPCLS primer and probe sequences

#### Primers:

| | |
|---|---|
| **ACTb_F** | CCTTTGCCGATCCGCCG |
| **ACTb_R** | CACGATGGAGGGGAAGACG |
| **COL1A1-F** | CAAGAGGAAGGCCAAGTCGAG |
| **COL1A1-R** | TTGTCGCAGACGCAGATCC |
| **ACTA2-F** | GCGTGGCTATTCCTTCGTTA |
| **ACTA2-R** | TTCTCAAGGGAGGATGAGGA |
| **FN1-F** | CCGACCAGAAGTTTGGGTTCT |
| **FN1-R** | ATGTCATGCTGCTTATCCCACT |

#### Probes:

| Oligo Name | Sequence (5' to 3') | 5' modification | 3' modification |
|---|---|---|---|
| COL1A1 _probe | GGT CAT GGT ACC TGA GGC CGT TCT GTA CG | VIC | MGBNFQ |
| ACTA2_probe | AGC GTG AGA TTG TCC GGG ACA TCA AGG A | ABY | QSY |
| ACTb_probe | TGG ATG ATG ATA TCG CCG CGC TCG TCG T | JUN | QSY |
| FN1_probe | TGG CTG CCC ACG AGG AAA TCT GCA CAA | FAM | MGBNFQ |

### Figure S1: Cell viability following 72 hours of incubation with select hit compound

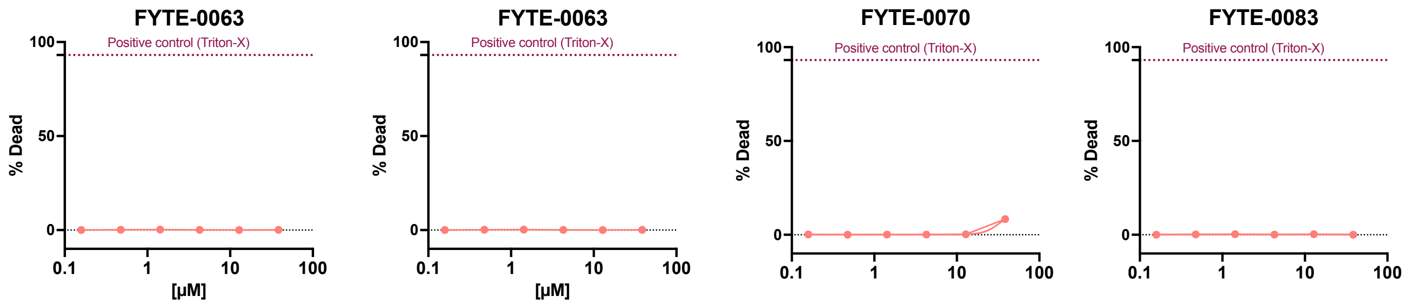


Following 72 hours of incubation with a dose range of hits, PI staining was used to calculate percent dead cells. In general, no loss of viability was observed at any tested dose.
